# SARS-CoV-2 drives NLRP3 inflammasome activation in human microglia through spike-ACE2 receptor interaction

**DOI:** 10.1101/2022.01.11.475947

**Authors:** Eduardo Albornoz, Alberto A Amarilla, Naphak Modhiran, Sandra Parker, Xaria X. Li, Danushka K. Wijesundara, Adriana Pliego Zamora, Christopher LD McMillan, Benjamin Liang, Nias Y.G. Peng, Julian D.J. Sng, Fatema Tuj Saima, Devina Paramitha, Rhys Parry, Michael S. Avumegah, Ariel Isaacs, Martin Lo, Zaray Miranda-Chacon, Daniella Bradshaw, Constanza Salinas-Rebolledo, Niwanthi W. Rajapakse, Trent Munro, Alejandro Rojas-Fernandez, Paul R. Young, Katryn J Stacey, Alexander A. Khromykh, Keith J. Chappell, Daniel Watterson, Trent M. Woodruff

## Abstract

Coronavirus disease-2019 (COVID-19) is primarily a respiratory disease, however, an increasing number of reports indicate that SARS-CoV-2 infection can also cause severe neurological manifestations, including precipitating cases of probable Parkinson’s disease. As microglial NLRP3 inflammasome activation is a major driver of neurodegeneration, here we interrogated whether SARS-CoV-2 can promote microglial NLRP3 inflammasome activation utilising a model of human monocyte-derived microglia. We identified that SARS-CoV-2 isolates can bind and enter microglia, triggering inflammasome activation in the absence of viral replication. Mechanistically, microglial NLRP3 could be both primed and activated with SARS-CoV-2 spike glycoprotein in a NF-κB and ACE2-dependent manner. Notably, virus- and spike protein-mediated inflammasome activation in microglia was significantly enhanced in the presence of α-synuclein fibrils, which was entirely ablated by NLRP3-inhibition. These results support a possible mechanism of microglia activation by SARS-CoV-2, which could explain the increased vulnerability to developing neurological symptoms akin to Parkinson’s disease in certain COVID-19 infected individuals, and a potential therapeutic avenue for intervention.

**SIGNIFICANCE STATEMENT:** Severe acute respiratory syndrome coronavirus 2 (SARS-CoV-2) principally affects the lungs, however there is evidence that the virus can also reach the brain and lead to chronic neurological symptoms. In this study, we examined the interaction SARS-CoV-2 with brain immune cells, by using an ex-vivo model of human monocyte-derived microglia. We identified robust activation of the innate immune sensor complex, NLRP3 inflammasome, in cells exposed to SARS-CoV-2. This was dependent on spike protein-ACE2 receptor interaction and was potentiated in the presence of α-synuclein. We therefore identify a possible mechanism for SARS-CoV-2 and increased vulnerability to developing neurological dysfunction. These findings support a potential therapeutic avenue for treatment of SARS-CoV-2 driven neurological manifestations, through use of NLRP3 inflammasome or ACE2 inhibitors.

## INTRODUCTION

Neuroinflammation is a hallmark of neurodegenerative diseases. A variety of stimuli within the central nervous system (CNS), including pathogens, injury, toxic metabolites, and protein aggregates among others, can lead to the activation of the innate immune response mainly through microglial activation. When chronically activated, this defence mechanism creates a proinflammatory environment that drives neurodegeneration (1, 2). Microglia are resident populations of macrophages in the CNS that respond to pathogen-associated molecular patterns (PAMPs) and host- or environment-derived danger-associated molecular patterns (DAMPs) to drive innate immune responses and inflammation within the brain. Recent evidence has highlighted the role of intracellular protein complexes, known as the inflammasomes, in CNS innate immunity.

These complexes mediate the response to PAMPs and DAMPs, leading to the generation of IL-1β, IL-18 and ultimately cellular pyroptosis, which can aid in the elimination of invading pathogens, clearance of damaged cells, and promotion of tissue repair (3). The NLR family pyrin domain containing 3 (NLRP3) inflammasome is a key inflammasome expressed by microglia (4), and is activated by multiple protein aggregates associated with neurodegenerative disease including α-synuclein in Parkinson’s disease (PD), amyloid-β in Alzheimer's disease (AD), and TDP43 and SOD1 aggregates in amyotrophic lateral sclerosis (ALS) (5–7). Microglial NLRP3 inflammasome can also be activated by a variety of pathogenic viruses with neurotropism such as Zika virus (ZIKV) and Japanese Encephalitis virus (JEV) (8, 9). The NLRP3 inflammasome is comprised of the NLRP3 protein, the adaptor molecule apoptosis-associated speck-like protein containing a CARD (ASC), and caspase-1. Activation of NLRP3 is a two-step process; a priming step usually mediated through a Toll-like receptor involves NF-κB-dependent induction of both NLRP3 and pro-IL-1β, whereas the triggering step leads to oligomerisation of NLRP3, recruitment of ASC, and recruitment and activation of caspase-1. Active caspase-1 then cleaves pro-IL-1β and pro-IL-18 into their active forms, and initiates pyroptotic cell death (10).

The hypothesis that viral infections can accelerate neurodegeneration is gaining attention with relevance to the current COVID-19 pandemic (11, 12). It has become clear that severe acute respiratory syndrome coronavirus 2 (SARS-CoV-2) can invade and affect multiple organs and tissues including the brain (13, 14). Post-mortem analysis of brains obtained from deceased SARS-CoV-2 patients showed extensive microglial activation with pronounced neuroinflammation in the brainstem (15, 16).

Moreover, accumulating evidence shows that acute and sub-acute neurological complications of SARS-CoV-2 infections are reported in up to 85% of patients not only with severe COVID-19, but also in mildly symptomatic or asymptomatic patients (17, 18). These manifestations include headache, dizziness, impaired consciousness, encephalopathy, delirium, confusion, seizure, gait difficulties, cerebrovascular events, and post-infectious autoimmunity (19). Peripheral disorders include Guillain-Barre-syndrome, myositis-like muscle injury, and notably, up to 65% of COVID-19 affected patients reported decreased sense of smell or hyposmia (18), which also is a common pre-motor symptom in PD (20). Additionally, reported cases of PD linked to COVID-19 (21–23), have triggered attention to evaluating SARS-CoV-2 infections and their impact on PD (24, 25). However, the specific mechanism of how SARS-CoV-2 could increase the risk of developing neurological manifestations, and potentially PD, and how this infection could possibly impact synucleinopathy has not been demonstrated.

Here, we used a human monocyte-derived microglia (MDMi) cellular model to assess NLRP3 inflammasome activation in response to SARS-CoV-2, and its spike protein, and the consequences of this exposure in the presence of α-synuclein protein aggregate fibrils. We determined that SARS-CoV-2 isolates, as well as spike protein alone, can both prime and activate the NLRP3 inflammasome in human microglia through NF-κB and ACE2. Microglia exposed to SARS-CoV-2, or its spike protein also potentiated α-synuclein mediated NLRP3 activation, indicating a possible mechanism for COVID-19 and increased vulnerability to developing movement disorders in certain infected individuals.

## RESULTS

### SARS-CoV-2 can enter human monocyte-derived microglia (MDMi) without supporting replication

To investigate the possible role of SARS-CoV-2 in promoting inflammasome activation in the brain, we first generated MDMi following an established protocol to obtain adult microglia (26). Initially, we verified that our generated MDMi highly expressed the typical microglia signature markers – P2RY12 and TMEM119 compared to monocyte derived macrophages (MDM) (Figure 1A-B). Next, we assessed whether these MDMi can support SARS-CoV-2 replication by monitoring infectious virus particle release after infection with multiplicity of infection (MOI) of 1 and 0.1 To achieve this, we used an early clinical isolate of SARS-CoV-2, referred to here as D614 (27). Notably, we found no secreted virus in the supernatant of infected MDMi and mouse primary microglia (mMi) cell culture supernatants, which contrasted to that in Vero E6 and Caco2 cells (Figure 1C), supporting the notion that microglia cells do not support SARS-CoV-2 replication *in vitro*.

**Figure 1:**
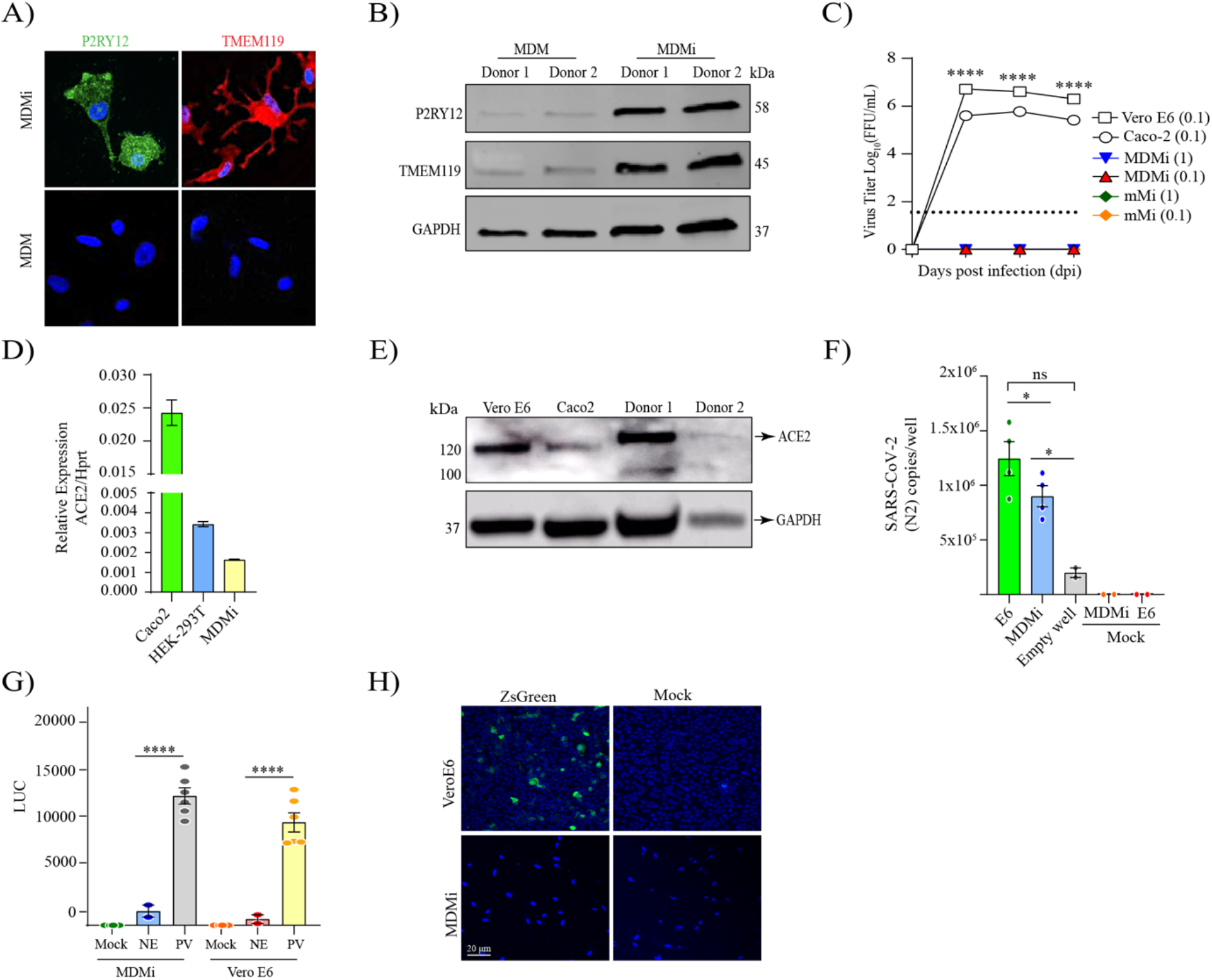
SARS-CoV-2 isolates can enter human monocyte-derived microglia (MDMi) in the absence of viral replication. Microglia signature markers, P2RY12 and TMEM119 (in green) and cells nuclei (in blue) assessed by immunofluorescence staining, and representative western blot are presented in panel (**A-B**), respectively. Growth kinetics of SARS-CoV-2 (D614) on MDMi, mouse microglia (mMi), Vero E6 and Caco2 cells in (**C**). Relative expression of ACE2 in MDMi by qPCR compared to Vero E6 and Hek-293T in (**D**). Level of ACE2 receptor in MDMi and mouse microglia compared to Caco2 and Vero E6 cells analysed by western blot shown in panel (**E**). Viral RNA levels from SARS-CoV-2 particles bound on cell surface expressed as N2 copies/well in (**F**) Intracellular luciferase level (LUC) delivered by pseudo-virus (PV) particle for SARS-CoV-2 in MDMi and Vero E6 compared to the non-glycoprotein control (NE) in (**G**). SARS-CoV-2 replication on MDMi (at MOI of 1) and Vero E6 (at MOI of 0.01) using SARS-CoV-2 reporter virus expressing ZsGreen fluorescent protein assessed directly under confocal microscopy at 3dpi are shown in panel (**H**). Data points are means + SEM from at least three different donors. *P < 0.05, **P < 0.01, and ***P < 0.001 and **** P < 0.0001 by two-way ANOVA test with Sidak’s correction.

As angiotensin-converting enzyme 2 (ACE2) is the most well-characterized receptor for SARS-CoV-2 cell attachment (28), and is expressed in the CNS, predominantly by glia, neurons and neurovascular endothelium (29), we proceeded to determine the level of ACE2 in MDMi by qPCR and western blot. We observed that MDMi expressed ACE2 mRNA, although the levels were lower compared to Vero E6 and Caco2 cells (Figure 1D). Western blot analysis from lysed microglia showed that ACE2 protein levels varied greatly in individual donors and displayed a differential pattern of expression compared to Vero E6 and Caco2 cells (Figure 1E). The control cells (VeroE6 and Caco2) showed an expected full-length size of the glycosylated ACE2 form of approximately 120 KDa, while microglia cells showed molecular weights of ~135 and ~100 kDa. Notably, similar patterns have been also found in endothelial cells and heart tissue from COVID-19 patients (30, 31).

Given that MDMi expressed ACE2 receptor, yet did not support virus replication, we sought to determine whether SARS-CoV-2 binds to the microglial cell surface. To address this, MDMi were exposed to D614 at MOI of 1, incubated for 2 hours at 4°C and then subjected to several washes to remove all unbound particles. We found significant levels of viral RNA from bound virus particle on the cell surface (Figure 1F), suggesting that SARS-CoV-2 can indeed bind to MDMi. We next investigated whether virus binding could promote viral entry by using a pseudo-virus entry assay as previously reported for SARS-CoV-2 (32, 33). MDMi were transduced with SARS-CoV-2 pseudo-virus for 72 hours and titer was determined by luciferase activity. We observed higher levels of intracellular luciferase activity in microglia cells infected with the pseudo-virus compared to the non-glycoprotein control (NE) and this level was comparable to pseudo-virus transduced VeroE6 cells (Figure 1G), suggesting that SARS-CoV-2 has the ability to enter human microglia cells.

To further investigate whether SARS-CoV-2 productively replicates in MDMi, we utilised a recently characterised SARS-CoV-2 reporter virus bearing ZsGreen fluorescent protein (34), and infected cells using a MOI of 1 and monitored up to 3 days post infection (dpi). As expected, we detected a high level of ZsGreen fluorescent protein expression in Vero E6 cells (Figure 1H and supplementary Figure 1). Furthermore, no intracellular ZsGreen fluorescence was observed in MDMi (Figure 1H and supplementary Figure 1), confirming the inability of SARS-CoV-2 to establish replication in these cells.

### The inflammasome is activated in human microglia by SARS-CoV-2

Microglia are the resident immune cells found in the CNS that patrol the brain sensing for pathogens or damage-associated stress signal (35). To investigate microglial inflammasome activation in response to SARS-CoV-2, we exposed MDMi cells to SARS-CoV-2 and measured key inflammasome activation signals. MDMi were incubated with MOI of 1 of D614 isolate directly, or in LPS-primed cells. At 24-hour post infection, western blot and immunocytochemistry was performed to examine markers of inflammasome activation (Figure 2A-C).

**Figure 2.**
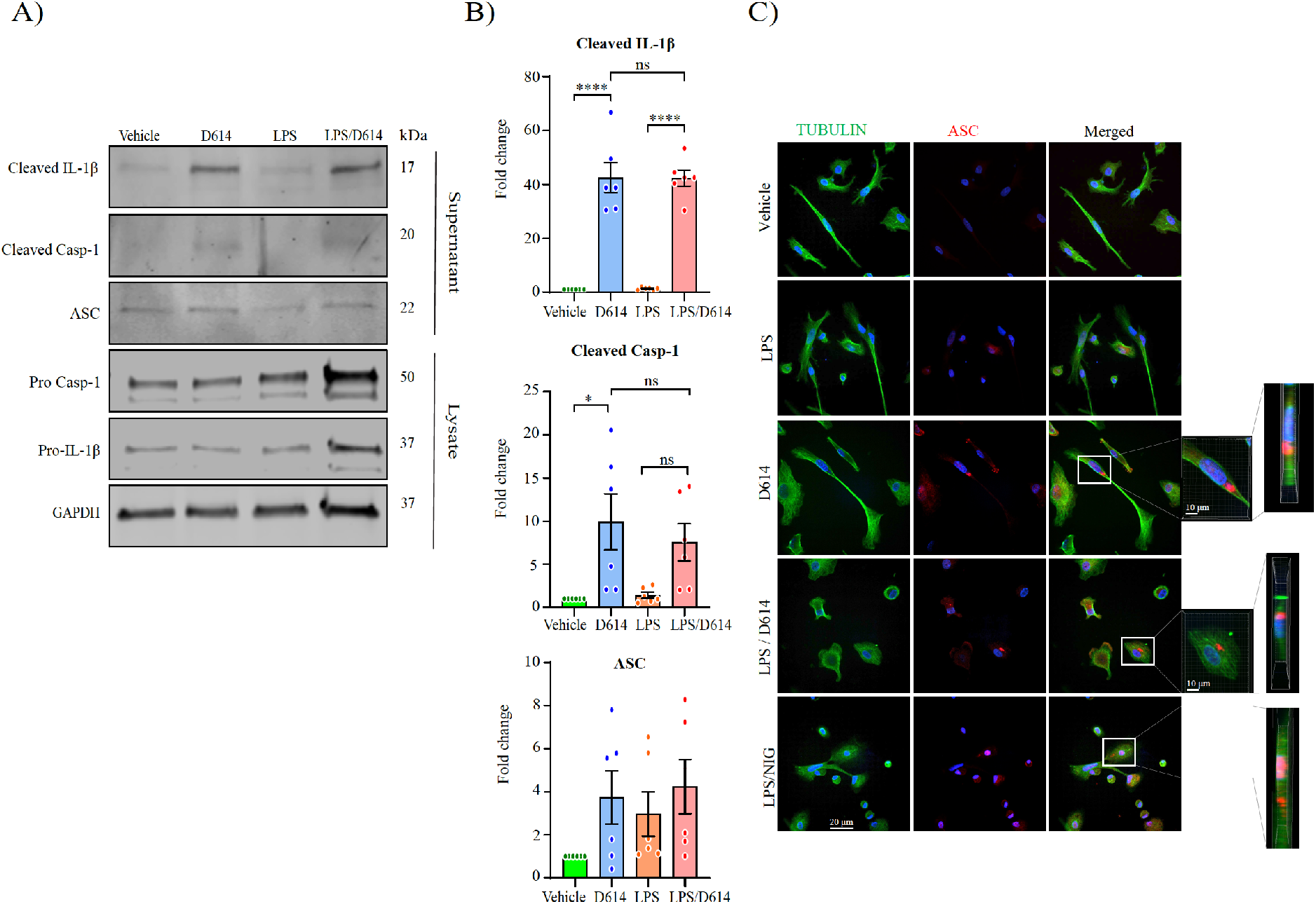
SARS-CoV-2 activates the inflammasome in MDMi. Western blot and densitometric analysis (fold change against vehicle group) for cleaved IL-1β, cleaved caspase-1 (p20), and ASC in the supernatants of LPS-primed or unprimed MDMi treated with SARS-CoV-2 (D614) isolate for 24 hours are presented in (**A-B**) respectively. Expression of GAPDH was determined in cell lysates. Immunofluorescence staining of primed or unprimed MDMi treated with SARS-CoV-2 (D614)–for Tubulin is stained in green and the formation of a characteristic inflammasome ASC speck (red) is shown in (**C**). LPS-Nigericin (Nig; 10μM, 1 hour) was used as a positive control. Scale bar, 20 μm. Inset magnified view of ASC specks. DAPI (blue), 4′,6-diamidino-2-phenylindole). Data points are means + SEM from at least three different donors. *P < 0.05, **P < 0.01, and ***P < 0.001 and **** P < 0.0001 by one-way analysis of variance (ANOVA) with Tukey’s post hoc test.

We identified SARS-CoV-2 alone induced inflammasome activation in MDMi as measured by the release of cleaved IL-1β in the supernatant of cells exposed to D614. This correlated with increased levels of cleaved caspase-1, validating activation of the inflammasome (Figure 2A-B). These results were corroborated by ASC speck formation, a cellular hallmark of inflammasome activation. Increased ASC speck staining was observed in MDMi cells treated with D614 (both primed and unprimed), and LPS-primed cells activated with nigericin (Nig) as a positive control (Figure 2C). Notably, our finding that SARS-CoV-2 exposure can directly activate the inflammasome in MDMi in the absence of priming, indicates that the virus can both prime and activate the inflammasome.

### SARS-CoV-2 spike protein activates the NLRP3 inflammasome in human microglia

Given that MDMi cells express ACE2 and that live virus activated MDMi independent of viral replication, we next determined whether spike protein itself could trigger inflammasome activation directly in human microglia. We previously designed a prefusion-stabilized SARS-CoV-2 spike protein (S-clamp) that resembles a closed trimeric prefusion conformation (36). To further identify the mechanism of inflammasome activation by SARS-CoV-2, we first produced low endotoxin S-clamp and a control trimeric fusion protein (F-clamp) from Nipah virus and validated these proteins using SDS-PAGE, size exclusion chromatography and ELISA (Figure 3A-C). As expected, the monomeric molecular weight of the S-clamp and F-clamp monomers were 180 and 60 KDa, respectively (Figure 3A). Additionally, we also confirmed that the majority of the S-clamp and F-clamp were presented in their trimeric form using size-exclusion chromatography and maintained reactivity assessed by binding of key specific antibodies (Figure 3B and C). Next, we proceeded to expose LPS-primed MDMi with different concentrations of S-clamp or control F-clamp protein as shown in a schematic representation in Figure 3D and identified that spike protein (S-clamp), but not the control F-clamp, induced significantly increased levels of IL-1β in supernatants after 24 hours exposure (Figure 3E). This activation was entirely ablated in the presence of MCC950, a selective inhibitor of NLRP3, confirming that the spike protein of SARS-CoV-2 is able to activate the NLRP3 inflammasome in LPS-primed microglia (Figure 3E). We further confirmed spike-mediated inflammasome activation through western blotting, showing dose-dependent increases in cleaved IL-1β and cleaved caspase-1 in the supernatant, and NLRP3 in cell lysates (Figure 3F-G). Although NLRP3 and pro-IL-1β are induced by LPS, they had evidently decayed after the wash-out of LPS but been re-induced by the spike protein. This this suggests that spike protein is both priming and activating the inflammasome.

**Figure 3.**
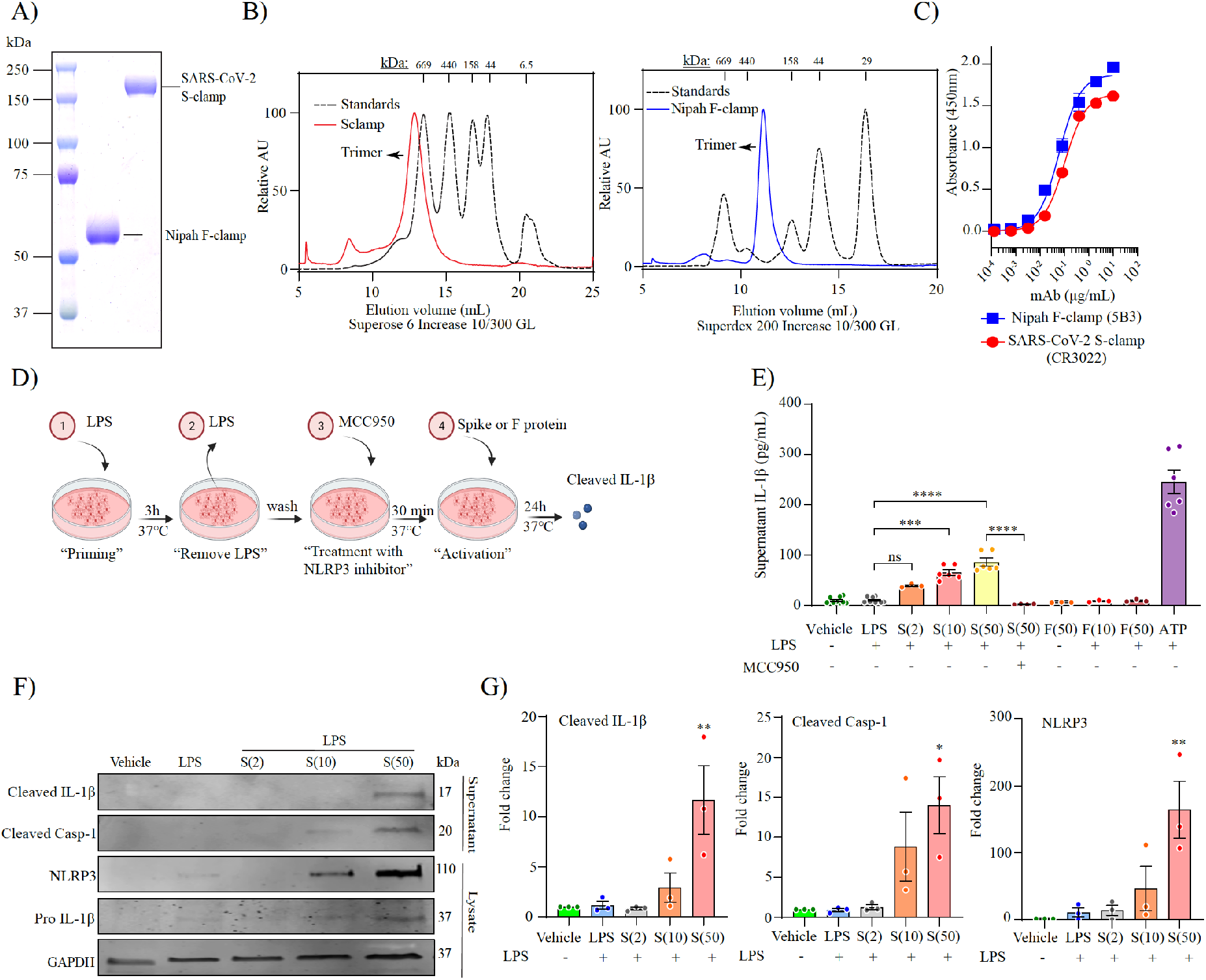
SARS-CoV-2 spike protein activates the NLRP3 inflammasome in LPS-primed MDMi. Prefusion-stabilized SARS-CoV-2 spike protein (S-clamp) and Fusion protein of Nipah virus (F-clamp) characterization by SDS-PAGE (**A**), size-exclusion high-performance liquid chromatography (**B**) and ELISA with conformational specific monoclonal antibodies (**C**). Schematic representation for spike activation on LPS primed-MDMi (D). Spike–mediated microglial IL-1β secretion (supernatant) in vehicle (untreated) or LPS-primed MDMi exposed to S-clamp (S; 2-50 μg) or F-clamp (F; 50 μg) in presence or absence of MCC950 (10 μM) treatment is shown in panel (**E**). ATP (5 mM) treatment for 1 hour was used as a positive control. Western blots (**F**) and densitometric analysis (fold change against vehicle group) (**G**) for NLRP3 in cell lysates of S-clamp–activated MDMi. Data are means + SEM from at least three different donors. *P < 0.05, **P < 0.01, and ***P < 0.001 and **** P < 0.0001 by one-way analysis of variance (ANOVA) with Tukey’s post hoc test.

### Spike protein activates NLRP3 inflammasome through ACE2 in human microglia

As spike protein is the major surface glycoprotein of the SARS-CoV-2 viral particle and contains a receptor-binding domain (RBD) that recognises ACE2 (37), we hypothesized that inflammasome activation is mediated by ACE2-RBD interaction. To address this, we first produced a low endotoxin version of a soluble human ACE2 protein (hACE2-FcM) and validated it in a neutralization assay. We found that the soluble human ACE2 protein blocked SARS-CoV-2 entry into Vero E6 cells (Figure 4A) with 50% inhibitory concentration (IC50) of 39 μg/mL, compared to control protein (NCAM-FcM) produced similar manner as hACE2-FcM (Figure 4A). Based on this finding, we complexed the soluble ACE2 protein with the S-clamp at a molar ratio of 5:1, respectively, incubated at 37°C for 1 hour, and then used this complex to treat MDMi. We found a complete inhibition of IL-1β secretion in culture supernatants compared to S-clamp treatment alone (Figure 4B). Additionally, pre-treatment of MDMi with the ACE2 inhibitor, MLN-4760, also significantly reduced spike protein induced IL-1β release from LPS-primed MDMi (Figure 4C). Furthermore, to confirm that MDMi activation is ACE2 dependent, we used a well characterised monoclonal antibody (3E8), against human ACE2 (38). To test the effect of 3E8 in blocking cell activation by spike protein, we first produced a low endotoxin level of 3E8 and a control antibody CO5 (anti HA of influenza A), followed by validation with SDS-PAGE and ELISA (Figure 4D-E). Pre-treatment of LPS-primed MDMi with 3E8 specifically inhibited IL-1β secretion after activation with S-clamp compared to CO5 and nigericin (Figure 4F), suggesting that spike-ACE2 interaction specifically contributes to inflammasome activation in microglia.

**Figure 4.**
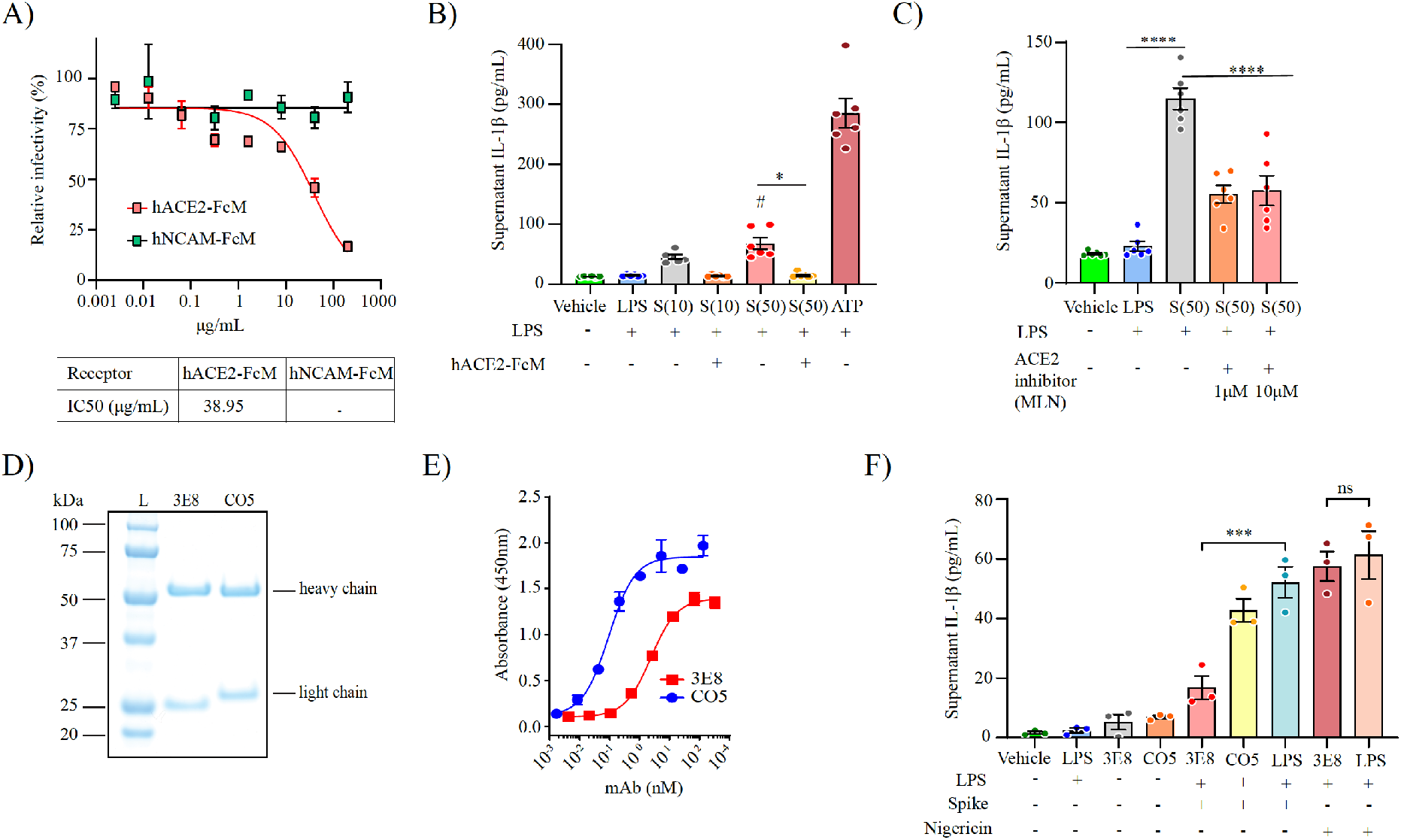
SARS-CoV-2 spike protein activates the NLRP3 inflammasome through ACE2 in MDMi. Relative of infectivity determined by Plaque Reduction Neutralisation Test (PRNT) to verify the neutralizing level of a soluble receptor hACE2-FcM compared to a non-related SARS-CoV-2 receptor NCAM-FcM (top) and the inhibitory concentration (IC50) (bottom) (**A**). Spike–mediated IL-1β secretion (supernatant) in vehicle (untreated) or LPS-primed MDMIs exposed to S-clamp (S; 10-50 μg) in presence or absence of the soluble hACE2-FcM protein. ATP (5 mM) treatment for 1 hour was used as a positive control (**B**). Inhibition of spike-mediated IL-1β secretion by ACE2 inhibition with MLN-4760 (1 or 10 μM) (**C**). Validation of low endotoxin anti-ACE2 (3E8) and anti-Hemagglutinin from influenza A H3 (CO5) proteins by SDS-PAGE and ELISA (**D-E**). Effect of 3E8 in blocking cells activation by spike protein in pre-treatment of LPS-primed MDMi exposed to S-clamp (**F**). Data are means + SEM from at least three different donors. *P < 0.05, **P < 0.01, and ***P < 0.001 and **** P < 0.0001 by one-way analysis of variance (ANOVA) with Tukey’s post hoc test.

### SARS-CoV-2 spike protein primes the NLRP3 inflammasome through NF-κB

We next evaluated if spike protein can also prime MDMi. To achieve this, MDMi were first stimulated with S-clamp for 6 hours, then media containing S-clamp was removed and replaced with fresh media and incubated with either ATP or nigericin for 1 hour as the inflammasome-activating signal, as shown in a schematic representation in Figure 5A. We observed a significant IL-1β release for both activators in S-clamp-primed cells, in comparison with either vehicle, S-clamp, ATP or nigericin alone, and the level was of a similar, but slightly reduced magnitude to LPS-primed cells (Figure 5B-C). We also confirmed an S-clamp priming effect through immunocytochemistry, showing ASC speck formation in S-clamp-primed cells activated with nigericin (Figure 5D), displaying a similar morphology to cells previously primed with LPS and activated with nigericin used as a positive control (Figure 2C).

**Figure 5.**
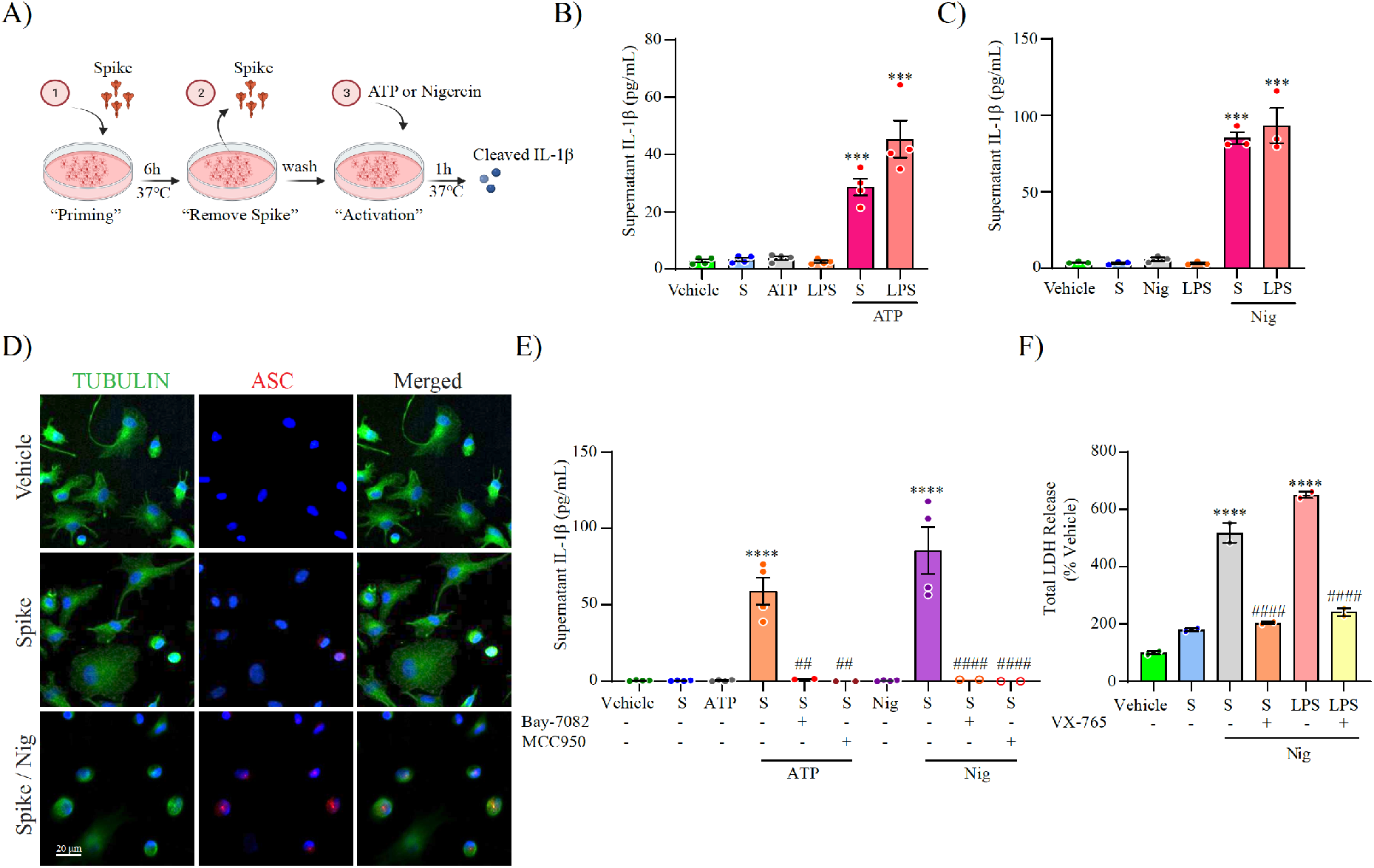
SARS-CoV-2 spike protein primes the NLRP3 inflammasome through NF-κB. Schematic representation for spike priming experiments (6 hours) followed ATP or Nigericin (Nig) activation (1 hour) (**A**). Level of secreted IL-1β in unprimed or S-clamp-primed MDMi (S; 50μg 6 hours) followed activation with ATP (5mM, 1 hour) in panel (**B**) or Nigericin (Nig; 10μM, 1 hour) in panel (**C**). In both LPS-primed cells were used as a positive control (200 ng/ml 3 hours). Immunofluorescence staining of vehicle or S-clamp (S; 50μg 6 hours)–primed MDMi, activated with Nigericin (Nig; 10μM, 1 hour) showing tubulin (green) and the formation of a characteristic inflammasome ASC speck (red) are shown in (**D**). Scale bar, 20 μm. ATP and Nigericin–mediated IL-1β secretion (supernatant) in vehicle (untreated) or S-clamp-primed MDMIs exposed to ATP (5mM, 1 hour) or Nigericin (Nig; 10μM, 1 hour) in presence or absence of Bay 11-7082 (3 μM) or MCC950 (10 μM) are shown in (**E**). Lactate dehydrogenase (LDH) release assay for quantification of caspase-1–dependent pyroptosis in S-clamp (S; 50 μg 6 hours) primed cells activated with Nigericin (Nig; 10μM, 1 hour) in (**F**). LPS-Nigericin and VX-765 (20 μM) were used as positive controls. Data are means + SEM from at least 3 independent donors. ***P < 0.001 and **** P < 0.0001 by one-way analysis of variance (ANOVA) with Tukey’s post hoc test.

To test if NF-κB signalling pathway is required for inflammasome priming by spike protein, we pre-treated MDMi with the NF-κB inhibitor, Bay 11-7082, before stimulation with S-clamp and the addition of ATP or nigericin. We found a complete inhibition of IL-1β release (Figure 5E), confirming S-clamp priming activity is mediated through the NF-κB pathway. Moreover, after priming with S-clamp, and before activating with either ATP or nigericin, we treated the cells with the NLRP3 inhibitor MCC950 and demonstrated a complete inhibition of IL-1β secretion (Figure 5E), confirming as expected that nigericin and ATP triggered NLRP3 after priming by spike protein.

Recently, it has been shown that SARS-CoV-2 triggers pyroptosis in human monocytes (39). Therefore, we assayed whether S-clamp priming and nigericin inflammasome activation triggered pyroptosis in MDMi. Pyroptosis was quantified using lactate dehydrogenase (LDH) release, and the caspase-1 inhibitor VX-765 was used to selectively assess the role for inflammasome activation in cell death. We observed that nigericin treatment of S-clamp primed MDMi readily triggered caspase-1-dependent pyroptosis within 1 hour (Figure 5F) which was significantly reduced in the presence of caspase-1 inhibitor VX-765. SARS-CoV-2 spike protein mediating priming of the inflammasome has recently be documented in macrophages derived from COVID-19 patients (40). Our data now provides strong evidence that SARS-CoV-2 spike protein can also prime the NLRP3 inflammasome through the NF-κB pathway in human microglia.

### SARS-CoV-2 promotes α-synuclein mediated NLRP3 inflammasome activation by priming MDMi through spike protein

The underlying mechanism of microglial activation by SARS-CoV-2 and their impact in presence of endogenous neurodegenerative disease-driving triggers are unclear. To better understand the effect of SARS-CoV-2 infection on promoting human microglia activation in relation to brain disease triggers, MDMi were incubated with D614 (MOI 0.1 or 1) in the presence or absence of preformed fibrils of α-synuclein for 24 hours as shown in a schematic representation in Figure 6A, and the level of cleaved IL-1β , cleaved caspase-1 and ASC in the supernatant were measured by western blot. We identified the presence of cleaved IL-1β, cleaved caspase-1, and ASC in the supernatant of MDMi treated with SARS-CoV-2 and α-synuclein, in the absence of LPS (Figure 6B). Notably, an increase of inflammasome activation was achieved with D614 (0.1 MOI) in presence of α-synuclein, whereas neither D614 at MOI 0.1, or α-synuclein alone, were able to activate the inflammasome; however, when combined, they released significant cleaved IL-1β, and cleaved caspase-1 in the supernatant (Figure 6B, C). Immunocytochemistry for ASC further confirmed this observation, showing increased levels ASC speck formation in cells treated with D614 at MOI 0.1 in presence of α-synuclein (Figure 6D).

**Figure 6.**
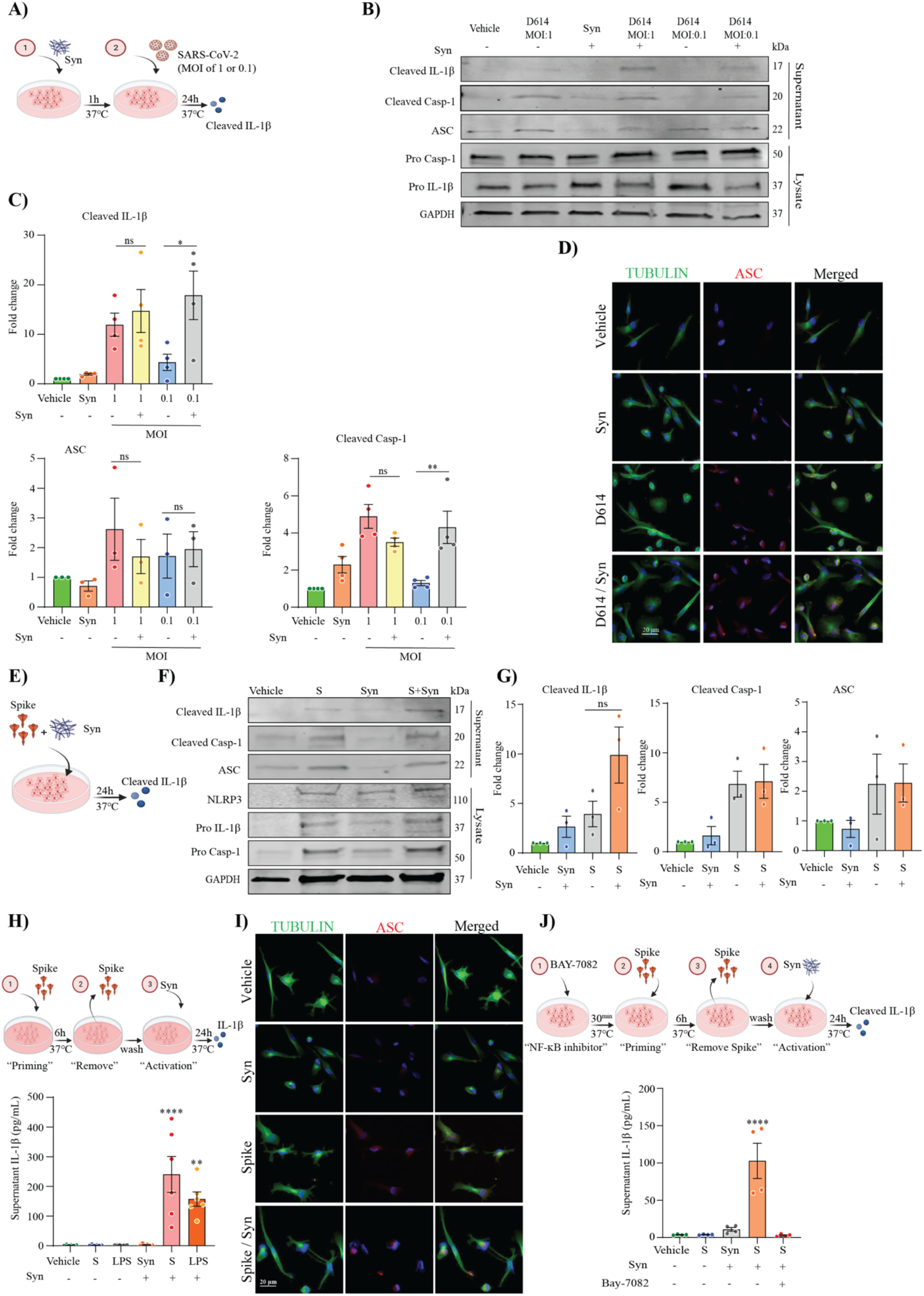
SARS-CoV-2 promotes α-synuclein mediated NLRP3 inflammasome activation, priming MDMi through spike protein. A schematic representation of SARS-CoV-2 exposure in presence of α-synuclein (Syn) for 24 hours on MDMi (**A**). Level of cleaved IL-1β, in the supernatants of MDMi, treated with either D614 (MOI 1, 0.1) or α-synuclein (Syn; 10 μM) or together for 24 hours by western blot and densitometric analysis are shown in panel (**B, C**), respectively. Expression of GAPDH was determined in cell lysates. Immunofluorescence staining of MDMi treated with either α-synuclein (Syn; 10 μM) or D614 (MOI 0.1) or together for 24 hours showing tubulin (green) and the formation of a characteristic inflammasome ASC speck (red) are shown in (**D**). Scale bar, 20 μm. A schematic representation of spike (S-clamp) exposure in presence of α-synuclein (Syn) for 24 hours on MDMi (**E**). Western blots and densitometric analysis (fold change against vehicle group) for cleaved caspase-1 (p20), cleaved IL-1β, and ASC in the supernatants of MDMi treated with either s-clamp (S; 50μg) or α-synuclein (Syn; 10 μM) or together for 24 hours are presented in panel (**F**) and (**G**). A Schematic representation for spike priming (6 hours) followed α-synuclein (Syn) activation for 24 hours in (**H**) and α-synuclein (Syn; 10 μM) –mediated IL-1β secretion (supernatant) in unprimed or S-clamp-primed MDMi (S; 50μg 6 hours) is shown in panel **(I)**. LPS-primed cells were used as a positive control (200 ng/ml 3 hours). Representative immunofluorescence of S-clamp (spike; 50μg 6 hours) primed MDMI activated with α-synuclein (Syn; 10 μM) for 24 hours showing staining for Tubulin (green) and ASC speck formation (red) in panel (**J**). A schematic representation for NF-κB inhibition on spike-primed MDMi (6 hours) followed α-synuclein (Syn) activation for 24 hours (**K**). IL-1β secretion in vehicle (untreated) or S-clamp-primed MDMIs exposed to α-synuclein (Syn; 10 μM) in presence or absence of Bay11-7082(3 μM) for 24 hours in panel (**L**). Data are means + SEM from at least three different donors. *P < 0.05, **P < 0.01 and **** P < 0.0001 by one-way analysis of variance (ANOVA) with Tukey’s post hoc test.

To examine the role of SARS-CoV2 spike protein in the context of α-synuclein microglial inflammasome activation, we repeated the experimental paradigm with spike in presence of α-synuclein for 24 hours as shown in a schematic representation in Figure 6E. Performing western blot for cleaved IL-1β, cleaved caspase-1 and ASC on supernatants, we confirmed inflammasome activation with both spike and α-synuclein in MDMi, and a trend towards an increase in cleaved IL-1β when α-synuclein and spike were combined (Figure 6F-G). We next proceeded to evaluate whether spike protein could prime MDMi for enhanced inflammasome activation driven by α-synuclein. We therefore primed MDMi with S-clamp for 6 hours followed by α-synuclein activation for 24 hours (Figure 6H). We confirmed by ELISA that no significant IL-1β release was induced by either S-clamp priming or α-synuclein when administered alone, however robust IL-1β release was found in S-clamp primed cells in the presence of α-synuclein, at levels even greater than LPS-primed cells (Figure 6H). This result correlated with ASC speck formation in S-clamp primed cells activated with α-synuclein (Figure 6I). To confirm the S-clamp priming effect in this context was also mediated through NF-κB signalling, we pre-treated cells with the NF-κB inhibitor Bay 11-7082 and demonstrated a complete inhibition of cleaved IL-1β release (Figure 6J). Altogether, our data demonstrates that SARS-CoV-2, and spike protein, can both prime and activate the NLRP3 inflammasome in human microglia, also potentiating activation in the presence of α-synuclein, supporting a possible risk factor for COVID-19 in Parkinson’s disease and neurodegeneration.

## DISCUSSION

Several recent clinical studies have documented increased inflammasome activity in response to SARS-CoV-2 infection, leading to immune dysregulation that is associated with COVID-19 severity (41–43). In the periphery, it has been observed that monocytes from COVID-19 patients have increased caspase-1 activation, and this was correlated with higher levels of plasma IL-1β in critically ill patients (39, 44). In the CNS, SARS-CoV-2 mediated activation of the inflammasome in microglia has not previously been directly demonstrated, but there is increasing evidence for microglial activation in COVID-19 patient brains. For example, post-mortem brain analysis from COVID-19 deceased patients identified enlargement of microglial cell soma and thickening of processes detected by staining of microglial markers Iba1, and TMEM119 (15, 16). Further, in three recent post-mortem COVID-19 cases, SARS-CoV-2 nucleocapsid protein, ACE2, and NLRP3 were found together in microglia (45). Building on these observational studies, here we provide mechanistic insight into the molecular requirements of SARS-CoV-2 inflammasome activation in human microglial cells.

SARS-CoV-2 entry into host cells has been thoroughly described and is mediated by the binding of viral spike protein to the human receptor ACE2 (46, 47). Our results show that microglial cells express ACE2 receptor, and although the level is relatively low, SARS-CoV-2 is able to enter these cells but does not establish viral replication (Figure1). This has also been shown for human *in vitro* differentiated myeloid dendritic cells (mDC) as well as M1 and M2 macrophages, where in contrast to Vero E6 controls, no infectious virus production of SARS-CoV-2 is observed up to 48 hours after inoculation (48). Our data also aligns with the work of Yang et. al. where it was demonstrated that human pluripotent stem cell (hPSC)-derived microglia also express ACE2 receptor and are permissive to SARS-CoV-2-pseudo-virus entry (33). Additionally, this study also found low or undetectable levels of viral RNA in hPSC-derived microglia exposed to infectious SARS-CoV-2, offering further evidence that while ACE2 mediated viral uptake is possible, hPSC-derived microglia do not support SARS-CoV-2 replication (33).

We also show that SARS-CoV-2 can activate the inflammasome in human microglia, through the read-out of cleaved IL-1β, cleaved caspase-1, and ASC speck formation in the supernatant (Figure 2). As it has been previously demonstrated that the interaction between ACE2 receptor and spike protein can induce the hyperactivation of NLRP3 in endothelial cells (49), we investigated the role of spike-ACE2 interaction relative to NLRP3 activation in microglia using a prefusion-stabilized SARS-CoV-2 spike protein (S-clamp) (36). To confirm that NLRP3 activation on MDMi was ACE2 dependent we used a soluble human ACE2 receptor (hACE2-FcM), an ACE2 inhibitor (MLN-4760), and a well characterized monoclonal antibody (3E8) (38). All three approaches confirmed that spike protein can activate NLRP3 in human microglia-like cells through ACE2.

Although we demonstrated that spike protein can activate the NLRP3 inflammasome in human microglia, it is worth noting that SARS-CoV-2 also encodes other viral proteins that could be involved in inflammasome activation. SARS-CoV-2 is comprised of a nucleocapsid protein (N), spike protein (S), membrane protein (M), and envelope protein (E), in addition to a series of accessory proteins (ORF3a, ORF6, ORF7a, ORF7b, ORF8, and ORF10). Previous studies with the original SARS coronavirus, SARS-CoV have shown that protein E and ORF3a activate NLRP3, forming multimeric complexes that act as ion channels activating the NLRP3 inflammasome with IL-1β release, driven through NF-κB (50–52). Moreover, recent evidence demonstrated that N-protein interacts directly with NLRP3, promoting the binding of NLRP3 with ASC, facilitating NLRP3 inflammasome assembly indicating another distinct mechanism of direct inflammasome activation through interaction of a viral protein with NLRP3 (53). Our findings now provide further information that SARS-CoV-2 spike protein contributes directly to activating the NLRP3 inflammasome through ACE2.

Priming of the inflammasome in cells is process necessary to induce transcriptional up-regulation of NLRP3 and pro-IL-1β (54). Our initial observation that the SARS-CoV-2 virus itself can trigger inflammasome activation in MDMi without the need for priming supports a role for vigorous virus-mediated inflammasome activation *in vivo*. We also confirmed that spike protein alone can prime the inflammasome through NF-κB in MDMi, allowing for NLRP3 activation with classical inflammasome activators ATP and nigericin, as has been previously reported in human monocytes, macrophages, and human lung epithelial cells (55, 56). These findings support that human coronavirus spike protein can induce innate immune responses through NF-κB signalling.

We previously documented that activation of microglial NLRP3 inflammasomes through α-synuclein fibrils is a major driver of dopaminergic neuronal loss in experimental PD (5). The accumulation of α-synuclein aggregates, as seen in Lewy bodies, and their spread throughout the brain is correlated with the stages of PD progression (57). Of importance to the present study, there are increasing reports of significant neurological complications from SARS-CoV-2 infection in human patients (15–17, 21, 58, 59). The correlation between viral infection and the manifestation of Parkinson-like symptoms has been described for a variety of viruses including influenza virus, Japanese encephalitis virus (JEV), and West Nile virus (WNV) infection resulting in tremor, myoclonus, rigidity, bradykinesia, and postural instability (60). Moreover, post-mortem analysis performed on WNV-infected individuals showed an increased level of α-synuclein (61). This finding prompted the hypothesis that α-synuclein is upregulated during infection as an antiviral factor in neurons, where it is proposed to act as a natural antimicrobial peptide to restrict viral infection in the brain (61, 62). However, a recent study indicated that there were no alterations in α-synuclein levels in serum and CSF of COVID-19 patients with neurological symptoms (63). These findings suggest that the reported cases of parkinsonism after SARS-CoV-2 infection could be a consequence of an increased proinflammatory environment, mediated by blood brain barrier (BBB) disruption (64), peripheral cell infiltration (65), and microglial activation (66). These processes could be enhanced in the presence of ongoing synucleinopathies, or risk factors such as aging and poor health, leading to an accelerated neuronal loss, correlated with the reported parkinsonism symptoms and possible susceptibilities to developing PD post-SARS-CoV-2 infection.

Here we addressed the impact of SARS-CoV-2 on microglia in presence of α-synuclein. We showed that SARS-CoV-2 promotes α-synuclein mediated NLRP3 inflammasome activation by priming MDMi through spike protein, providing *ex vivo* support for the negative impact of SARS-CoV-2 on neurodegenerative diseases such as PD. It is also worth noting that there are several lines of evidence in the literature indicating neurological complications resulting from SARS-CoV-2 infection. These include: *i*) neuroinvasion by SARS-CoV-2 is demonstrated in humans, macaques and mice overexpressing human ACE2 (15, 67–70); *ii*) an extended microglial activation with pronounced neuroinflammation is reported in brain autopsies obtained from deceased SARS-CoV-2 patients (15, 16, 71); *iii*) significant deterioration of motor performance and motor-related disability is seen in PD patients recovering from COVID-19 (72, 73); and *iv*) Lewy body formation occurs in SARS-CoV-2 infected macaques (70). Thus, our finding complements the knowledge-gap in molecular mechanisms by which SARS-CoV-2 may activate microglia and lead to neurological manifestations. Our data suggest that the spike protein-mediated priming and/or activation of microglia through the ACE2-NF-κB axis may promote NLRP3 inflammasome activation leading to neuroinflammation and neurological phenotypes. Further, this process may be enhanced in the presence of neurodegenerative disease triggers such as α-synuclein aggregates, supporting a possible role for COVID-19 in triggering brain diseases such as PD. Since NLRP3 inhibitors are currently in clinical development for neurodegenerative diseases, including PD (5, 74), these findings also support a potential therapeutic avenue for treatment of SARS-CoV-2 driven neurological manifestations.

## MATERIALS AND METHODS

### Study design

Studies were primarily designed (i) to determine whether increased NLRP3 inflammasome activation occurs in human monocyte-derived microglia (MDMi) exposed to SARS-CoV-2 isolates and (ii) to evaluate this activation in the presence of preformed fibrils of α-Synuclein (Syn).

### Ethics and biological safety

Ethical approval for collecting and utilising human donor blood was obtained from The University of Queensland Human Research Ethics Committee (HREC approval #2020000559). All experiments with pathogenic SARS-CoV-2 were conducted under a certified biosafety level-3 (BSL-3) conditions in the School of Chemistry and Molecular Biosciences at The University of Queensland (SCMB-UQ). All personnel used powered air purifying respirator (PAPR; SR500 Fan Unit) as a respiratory protection at all the time within the facility. Surface disinfection was performed using 70% ethanol, while liquid and solid waste were steam-sterilized by autoclave. This work was approved by the Institutional Biosafety Committee from The University of Queensland (UQ) (UQ IBC, approvals IBC/390B/SCMB2020, IBC/1301/SCMB/2020, IBC/376B/SBMS/2020 and IBC/447B/SCMB/2021).

### Cells lines

Vero E6 cells (African green monkey kidney cell clones) and Caco-2 (human colorectal adenocarcinoma cells), and human embryonic kidney 293T cells (HEK293T) were maintained in Dulbecco's Modified Eagle Medium (DMEM) supplemented with 10% heat inactivated foetal calf serum (FCS) (Bovogen, USA), penicillin (100 U/mL) and streptomycin (100 μg/mL) (P/S) and maintained at 37 °C with 5 % CO_2_. All cell lines were verified to be mycoplasma free by first culturing the cells in antibiotic-free media and then subjected to a mycoplasma tested using MycoAlert™ PLUS Mycoplasma Detection Kit (Lonza, UK).

### Generation of human Monocyte-Derived Microglia (MDMi)

Monocytes were isolated from healthy donor blood collected into lithium heparin vacutainer tubes (Becton Dickinson) by a qualified phlebotomist, or from buffy coats obtained from Australian Red Cross Lifeblood as previously described (75). Briefly, donor blood or buffy coat was diluted 1:1 with phosphate buffered saline (PBS) and transferred into sterile SepMate 50 (STEMCELL Technologies, BC, Canada) as per manufacturer’s instructions. Peripheral blood monocytes (PBMCs) were collected. Monocytes were positively selected from whole PBMCs using anti CD14^+^ microbeads (Miltenyi Biotec) and plated at the following densities per well: 1 × 10^5^ cells (96-well plate) and 3 × 10^5^ cells (24-well plate). To induce the differentiation of MDMi, we incubated monocytes under serum-free conditions using RPMI-1640 Glutamax (Life Technologies) with 1% penicillin/ streptomycin (Lonza) and Fungizone (2.5 μg/ml; Life Technologies) and a mixture of the following human recombinant cytokines: M-CSF (10 ng/ml; Preprotech, 300-25), GM-CSF (10 ng/ml; Preprotech,300-03), NGF-β (10 ng/ml; Preprotech, 450-01), MCP-1(CCL2) (100 ng/ml; Preprotech, 300-04), and IL-34 (100 ng/ml; Preprotech, 200-34-250) under standard humidified culture conditions (37°C, 5% CO_2_) for up to 14 days. Differentiation of PBMCs into MDMi was confirmed by western blot and immunofluorescence for microglial markers compared to monocyte-derived macrophages (MDM).

### MDMi Treatments

For inflammasome activation experiments, MDMi were primed with 200 ng/ml of ultrapure LPS (*E.Coli* 0111:B4, Invivogen) for 3 hours or 50 μg of S-clamp or F-clamp for 6 hours. Cells were washed in after priming to remove residual LPS or S-Clamp and cells were stimulated with conventional NLRP3 inflammasome activators ATP (5 mM, Sigma) and nigericin (10 μM, Invivogen), or fibrillar α-synuclein (10 μM, Proteos), S-Clamp (2-50 μg) or SARS-CoV-2 isolates (MOI 0.1, 1) for the indicated time. For priming studies MDMis were pre-treated with the NF-κB inhibitor, Bay 11-7082 (3 μM, Sigma), before stimulation with S-clamp and the addition of ATP, nigericin or α-synuclein. For inhibition studies, MCC950 (10 μM), VX-765 (20μM, Invivogen) and MLN-4760 (1,10 μM, Sigma) were added after the priming step. At the end of treatment, the supernatant was collected and stored at −80°C until analysis by enzyme-linked immunosorbent assay (ELISA) or western blotting.

### Quantification of Caspase-1 Mediated Pyroptosis

At the end of each treatment, supernatants were collected and LDH release was quantified using an LDH assay kit (TOX7-Sigma) as per the manufacturer’s instructions. Caspase-1 dependent LDH release that was inhibited by the caspase-1 inhibitor VX-765 (20 μM), was used as a readout for pyroptosis as previously described (76).

### Viral isolate

SARS-CoV-2 were isolated from patient nasopharyngeal aspirates via inoculation in Vero E6 cells. An early Australian isolate hCoV-19/Australia/QLD02/2020 (QLD02) (GISAID Accession ID; EPI_ISL_407896) sampled on 30/01/2020 and named in this study as D614. This virus isolated was provided by Queensland Health Forensic & Scientific Services, Queensland Department of Health as passage 2 in Vero E6 cells. Viral stocks (passage 3) were then generated on VeroE6 cells and stored at −80 °C. To ensure there was no passage-to-passage variation of viruses used in this study or loss of the spike furin cleavage on VeroE6 passaged SARS-CoV-2 isolate whole SARS-CoV-2 sequencing and variant bioinformatics analysis was conducted as per (34) to QLD02 isolate. Briefly, the nCoV-2019 Nanopore sequencing protocol v3 (Josh Quick, University of Birmingham) was used with minor modifications. RNA was isolated from cell culture supernatant, and cDNA generated using Protoscript II first-strand cDNA synthesis kit as per manufacturer’s protocol (New England Biolabs, USA). SARS-CoV-2 cDNA was subsequently amplified using ARTIC network v2 primers using two-step PCR amplification with Q5® High-Fidelity DNA Polymerase (New England Biolabs, USA). PCR fragments were purified using AMPure XP beads (Beckman Coulter, USA) and subjected to End Repair/dA-Tailing using the NEBNext® Ultra™ II Module (New England Biolabs, USA). Passage 3 of QLD02 sample was multiplexed using the Native Barcoding Expansion kit (EXP-NBD104, Oxford Nanopore, UK) and Ligation Sequencing Kit (SQK-LSK109, Oxford Nanopore, UK). Prepared libraries were then quantified and loaded into equimolar concentrations totalling 20 fmol into a Flongle flow cell (FLOFLGOP1, Oxford Nanopore, UK). Variant analysis was conducted using iVar (v1.2.2) (77) and depth of sequencing coverage and consensus positions were visualized and calculated using Integrative Genomics Viewer (Version: 2.7.0).Virus stock titre was determined by an immuno-plaque assay (iPA) as previously described (78).

### Growth Kinetics

SARS-CoV-2 (D614) replication kinetics were assessed on VeroE6, Caco3, mouse primary microglia and MDMi cells. Briefly, 5×10^5^ cells were seeded in 24-well plates one day before infection. Cells were infected at a MOI of 1 or 0.1 for 30 min at 37 °C. The monolayer was washed five times with 1mL of additive-free DMEM and finally incubated with 1 mL of DMEM (supplemented with 2% FCS and P/S) at 37 °C with 5% CO_2_. Infectious viral titres were assessed in samples harvested from supernatant at the time points; 0-, 1-, 2- and 3-days post-infection (dpi). The viral titre was determined by an immuno-plaque assay (iPA) on VeroE6 cells. Two independent experiments were performed with 2 technical replicates.

### Binding assay

Vero E6 and MDMi cells were inoculated with an MOI=1 of SARS-CoV-2 (D614) for 2 hours at 4 °C and then, cells were washed 8 times with fresh cold media to remove unbound viruses. Cells were then harvested in TRI Reagent (Millipore, Sigma-Aldrich, Germany) and a total RNA was purified and quantified by a real time RT-PCR.

### RNA extraction

RNA was extracted using TRIzol (Thermo Fisher) plus an RNA extraction RNeasy Micro kit (Qiagen). Briefly, the aqueous phase containing the RNA was separated by adding chloroform to the TRIzol samples and centrifuging them for 30 min, 4°C at 12,000 g. An equal volume of 70% ethanol was added to the isolated aqueous phase, mixed and then added to the RNeasy MinElute spin column. The following washes, DNase I treatment and elute steps were performed as described in the Qiagen RNeasy Micro Handbook. RNA samples were eluted in RNase free water.

### Quantitative Real-time PCR

One-step quantitative real-time PCR was performed in a QuantStudio 6 (Thermo Fisher) using GoTaq® Probe 1-Step RT-qPCR System (Promega). The CDC SARS-CoV-2 nucleoprotein N2 primer set was used for amplification. Forward: 5’-TTA CAA ACA TTG GCC GCA AA-3’, Reverse: 5’-GCG CGA CAT TCC GAA GAA-3’ and Probe: 5’-FAM-ACA ATT TGC CCC CAG CGC TTC AG-BHQ1-3’ (79). The standard curve was done using the 2019-nCoV_N Positive Control nucleoprotein DNA (IDT). A fixed volume representing 1/16 of the total RNA contained in each sample (well) was added to the master mix. Total number of copies were calculated using a semi-log line regression using GraphPad Prism 8.0.1. The human ACE2 transcript variant 2 was amplified using the OriGene primer set Forward: 5’-TCC ATT GGT CTT CTG TCA CCC G-3’ and Reverse: 5’-AGA CCA TCC ACC TCC ACT TCT C-3’. HPRT was use as the control housekeeping gene using primers, Forward: 5’-TCA GGC AGT ATA ATC CAA AGA TGG T-3’ and Reverse: 5’-AGT CTG GCT TAT ATC CAA CAC TTC G-3’. The relative expression is equal to the 2^(*−Δct*)^.

### Pseudo-virus entry assay

Pseudo-virus particles for SARS-CoV-2 were generated by using a lentiviral-based pseudo-particles system as previously described (32). Briefly, HEK293T cells were co-transfected with the following plasmids: 1 μg of p8.91 (encoding a second-generation lentiviral packaging plasmid for HIV-1 gag-pol), 1.5 μg of pCSFLW (firefly luciferase reporter gene) and, 1 μg of plasmid encoding SARS-CoV2 spike (D614) with C-terminal 18 amino acid deletion using Lipofectamine LTX Plus reagent (Invitrogen, USA) as per manufacturer’s protocol. A non-glycoprotein control (NE) was also generated using the same combination of plasmids as above, replacing plasmid encoding SARS-CoV2 spike, with 1 μg of an empty plasmid vector (pcDNA2.1).

Fourteen hours post-transfection, the medium was replaced with DMEM supplemented with 10% FCS and P/S and maintained at 37 °C with 5 % CO_2_ for 3 days. The supernatant containing SARS-CoV-2 pseudo-virus particles and the non-glycoprotein control (NE) was spun down at 3000 × g for 10 min at 4°C to remove cellular debris, aliquoted and stored at −80°C. To validate the pseudo-virus activity, approximately 2 × 10^4^ cells/well of HEK293T were seeded on a black flat-bottomed 96-well plate (Corning, USA) precoated with poly-L-lysine and incubated overnight at 37 °C with 5 % CO_2_. SARS-CoV-2 pseudo-virus particles and the non-glycoprotein control (NE) were diluted 1:5 and 1:20 in DMEM media supplemented with 10% FCS and P/S before infecting the cells with 100μL. HEK293T were incubated at 37 °C with 5 % CO_2_ for 3 days. The intracellular luciferase level was measured on Varioskan LUX multimode microplate reader (ThermoFisher, USA) by replacing the medium with 50 μL Bright-Glo substrate (Promega, USA), as per manufacturer’s protocol. The results showed a high level of intracellular luciferase in HEK293T compared to the non-glycoprotein control (NE) validating this batch of pseudo-virus particle for SARS-CoV-2 used in this study (Supplementary file 2). MDMi and Vero E6 were treated similarly with 1:5 dilution of Pseudo-virus particles stock and the level of intracellular luciferase was measured after 3 days.

### Plaque Reduction Neutralisation Test (PRNT)

The neutralisation levels of a soluble recombinant angiotensin-converting enzyme 2 (ACE2) receptor against the SARS-CoV-2 (D614) were verified by iPA (78). The generation of a soluble recombinant version of human ACE2 protein was recently described (36).

### Purification and analysis of Nipah F-clamp and SARS-CoV-2 S-clamp proteins

SARS-CoV-2, termed S-clamp, Nipah F-clamp (GenBank: NP_112026.1) and the soluble hACE2 proteins were expressed in ExpiCHO cells and purified as previously described (36, 80). Purified proteins were then characterised via SDS-PAGE, ELISA using ectodomain-specific monoclonal antibodies and size-exclusion chromatography. For SDS-PAGE, 4 μg of purified protein was mixed with DTT and analysed using a NuPAGE™ 4-12% Bis-Tris mini protein gel (ThermoFisher) as per the manufacturer’s instructions. Proteins were visualised by Coomassie staining.

For ELISA analysis, Nipah F-clamp, or SARS-CoV-2 S-clamp proteins were diluted to 2 μg/mL in PBS and coated overnight on Nunc MaxiSorp ELISA plates. The next day, plates were blocked with 150 μL/well of 5% KPL Milk Diluent/Blocking Solution Concentrate (SeraCare) in PBS with 0.05% Tween 20 (PBS.T) for 1 hour at room temperature. Blocking buffer was removed, and serial dilutions of Nipah F- or SARS-CoV-2 S ectodomain-specific antibodies, 5B3 and CR3022 (81), were added. Plates were incubated for 1 hour at 37 °C before they were washed three times with water and patted dry. Next, an HRP-conjugated goat anti-human secondary antibody (ThermoFisher) was added, and the plates incubated for 1 hour at 37 °C. The plates were washed and dried as above before the addition of TMB Single Solution chromogen/substrate (Invitrogen). Plates were allowed to develop for 5 minutes at room temperature before the reaction was stopped by the addition of 2N H2SO4. Absorbance at 450 nm was read on a Varioskan LUX Multimode Microplate Reader (ThermoFisher). Data were analysed using GraphPad Prism version 8 using a one site – specific binding model.

Nipah F-clamp and SARS-CoV-2 S-clamp proteins were further analysed for their oligomeric state via size-exclusion chromatography using a Superdex 200 Increase 10/300 GL or Superose 6 Increase 10/300 GL column, respectively. Approximately 30-50 μg of protein in PBS was loaded onto the column using an ÄKTA pure FPLC system at a flowrate of 0.5 mL/ minute. The limulus amebocyte lysate (LAL) based testing of endotoxin levels of the purified proteins were performed using Endosafe®-PTS™ device and cartridges according to the manufacturer’s protocol (Charles River). Additionally, low endotoxin levels for the soluble hACE2, and monoclonal antibodies (3E8 and CO5) were also produced and validated by SDS-PAGE and ELISA. For ELISAs, 2μg/mL in PBS of S-clamp or recombinant hACE2 or Hemagglutinin (HA) from influenza A H3 (Switzerland 2013) were coated overnight on Nunc MaxiSorp ELISA plates and then used to validate the hACE2, 3E8 and CO5 proteins respectively. The endotoxin levels were 1.1 EU/mg, 5.9 EU/mg, < 5 EU/mg, < 5 EU/mg and 124 EU/mg for SARS-CoV-2 S-clamp, Nipah F-clamp, hACE2, 3E8 and CO5 proteins, respectively.

### Preparation of fibrillar α-synuclein

Recombinant human α-synuclein monomer was obtained from Proteos, and *in vitro* fibril generation was performed with a final concentration of 2 mg/ml in phosphate-buffered saline (PBS) by incubation at 37⁰C with agitation in an orbital mixer (400 rpm) for 7 days with daily cycles of sonication used to break down fibrillar aggregates as outlined previously (5). The generation of fibrillar α-synuclein species was confirmed by transmission electron microscopy and Thioflavin T fluorescence prior to use.

### Western blotting

Primary microglial cells were collected and lysed using RIPA buffer (ThermoFisher). Proteins were separated in precast BioRad gradient (4-20%) gels. Proteins were then transferred to a nitrocellulose membrane and blocked for 1 h at room temperature (RT) using fluorescence western blocking buffer (Licor Bioscience). Membranes were washed 5 times with 5 min incubations per wash using either PBS containing 0.05% Tween-20 or tris(hydroxymethyl)aminomethane (Tris)-buffered saline containing 0.05% Tween-20. Primary human/rabbit/mouse antibodies, diluted in either Licor blocking buffer or 5% bovine serum albumin (BSA) solution, were then added to the membranes and incubated overnight at 4°C. Glyceraldehyde 3-phosphate dehydrogenase (GAPDH) was used as loading controls. Following 5 washes, respective infrared-dye or horseradish peroxidase (HRP) conjugated secondary antibodies against primary antibodies was added to the membranes for 1 hour at RT. Bands were visualised using either the Odyssey CLx imaging system (LI-COR) or enhanced chemiluminescence via SuperSignal^®^ West Pico Plus Chemiluminescent Substrate (Thermo-Scientific) accordingly to manufacturer’s instructions. Densitometric analysis was performed using Image Studio Lite software and the normalized band intensities were expressed as fold change over the control group.

### ELISA

Human IL-1 beta/IL-1F2 Duoset ELISA kit (R&D Systems, Catalog # DY201 was used to measure IL-1β in the supernatants of activated microglia. The assay was carried out according to the manufacturer’s instructions.

### MDMI Immunocytochemistry

MDMI cells were fixed with 4% paraformaldehyde in PBS for 10min. Cells were permeabilised with 0.1% triton X-100, subsequently washed three times in PBS, then blocked with 3% donkey serum in PBS for 1 hr at room temperature. Following this, cells were incubated for 2 hrs at room temperature with combinations of the following antibodies: rabbit anti-TMEM (1:100), rabbit anti-ASC (1:250), and mouse anti-Tubulin (1:250). Cells were then washed three times with PBS before incubating with secondary antibodies for 1 hr at room temperature. Secondary antibodies used were Donkey anti-rabbit 555 (1:2000) and donkey anti-mouse 488 (1:2000). Following three washes in PBS, cells were incubated with DAPI (1:4000 in PBS) before being mounted with Prolong gold antifade medium (Invitrogen) for fluorescent microscopy.

### Fluorescence microscopy methods

Images were collected using a Diskovery spinning disk confocal microscope (Andor/Nikon), 60XC CFI Plan Apochromat WI (NA 1.2) lens, with a disk pinhole size of 70um, and an Andor Zyla 4.2 sCMOS camera (Andor, UK). Images were collected at 12-bit and 2048 × 2048 pixel resolution. System settings, camera exposure times, and image brightness and contrast were consistent across all samples and optimised on Imaris 9.1.0 (Bitplane, UK) to create representative images for presentation. Samples were stained using the following fluorescent secondary antibodies: Donkey anti-rabbit Alexa Fluor® 555, Donkey anti-mouse Alexa Fluor® 488, and DAPI nuclear stain (brand). These fluorophores were then captured using laser lines 561nm, 488nm, and 405nm for excitation with appropriate filters to maximise emission photon capture. Images were captured in a Z-stack for intracellular ASC speck visualisation, with a step size of 0.15um.

### Statistical Analysis

All data were analysed using Prism software (GraphPad 9.1.2). For growth kinetics, a multiple comparison using two-way ANOVA test with Sidak’s correction was used to compare within groups. Statistical significance was set at 95 % (p = 0.05). For PRNT, a nonlinear regression with Inhibitor vs. response (three parameters) model was used to determine the best-fit curve. Data are represented as mean +/− s.e.m. from at least 3 experiments or donors. ANOVA followed by Tukey’s post-test was performed to compare all treatment groups in densitometries and ELISA. *P < 0.05, **P < 0.01, and ***P < 0.001 denote statistically significant differences between indicated groups.

## ACKNOWLEDGMENTS

We would like to thank the Queensland Health Forensic and Scientific Services, Queensland Department of Health, for providing the QLD02 SARS-CoV-2 isolate. We acknowledge funding support from the National Health and Medical Research Council (2009957), The Australian Infectious Diseases Research Centre (COVID-19 seed grant to AAK), and the Medical Research Future Fund (APP1202445-2020 MRFF Novel Coronavirus Vaccine Development Grant).

## Notes

### Competing Interest Statement

The authors have declared no competing interest.

## REFERENCES

1. E. A. Albornoz, T. M. Woodruff, R. Gordon, Inflammasomes in CNS Diseases. Exp Suppl 108, 41–60 (2018).

2. S. Voet, S. Srinivasan, M. Lamkanfi, G. van Loo, Inflammasomes in neuroinflammatory and neurodegenerative diseases. EMBO Mol Med 11 (2019).

3. F. Martinon, K. Burns, J. Tschopp, The inflammasome: a molecular platform triggering activation of inflammatory caspases and processing of proIL-beta. Mol Cell 10, 417–426 (2002).

4. J. G. Walsh, D. A. Muruve, C. Power, Inflammasomes in the CNS. Nat Rev Neurosci 15, 84–97 (2014).

5. R. Gordon et al., Inflammasome inhibition prevents alpha-synuclein pathology and dopaminergic neurodegeneration in mice. Sci Transl Med 10 (2018).

6. M. T. Heneka et al., NLRP3 is activated in Alzheimer’s disease and contributes to pathology in APP/PS1 mice. Nature 493, 674–678 (2013).

7. V. Deora et al., The microglial NLRP3 inflammasome is activated by amyotrophic lateral sclerosis proteins. Glia 68, 407–421 (2020).

8. D. K. Kaushik, M. Gupta, K. L. Kumawat, A. Basu, NLRP3 inflammasome: key mediator of neuroinflammation in murine Japanese encephalitis. PLoS One 7, e32270 (2012).

9. W. Wang et al., Zika virus infection induces host inflammatory responses by facilitating NLRP3 inflammasome assembly and interleukin-1beta secretion. Nat Commun 9, 106 (2018).

10. K. V. Swanson, M. Deng, J. P. Ting, The NLRP3 inflammasome: molecular activation and regulation to therapeutics. Nat Rev Immunol 19, 477–489 (2019).

11. G. De Chiara et al., Infectious agents and neurodegeneration. Mol Neurobiol 46, 614–638 (2012).

12. L. Ferini-Strambi, M. Salsone, COVID-19 and neurological disorders: are neurodegenerative or neuroimmunological diseases more vulnerable? J Neurol 268, 409–419 (2021).

13. J. Liu et al., SARS-CoV-2 cell tropism and multiorgan infection. Cell Discov 7, 17 (2021).

14. V. G. Puelles et al., Multiorgan and Renal Tropism of SARS-CoV-2. N Engl J Med 383, 590–592 (2020).

15. K. T. Thakur et al., COVID-19 neuropathology at Columbia University Irving Medical Center/New York Presbyterian Hospital. Brain 10.1093/brain/awab148 (2021).

16. J. Matschke et al., Neuropathology of patients with COVID-19 in Germany: a post-mortem case series. Lancet Neurol 19, 919–929 (2020).

17. M. A. Ellul et al., Neurological associations of COVID-19. Lancet Neurol 19, 767–783 (2020).

18. M. Merello, K. P. Bhatia, J. A. Obeso, SARS-CoV-2 and the risk of Parkinson’s disease: facts and fantasy. Lancet Neurol 20, 94–95 (2021).

19. S. Najjar et al., Central nervous system complications associated with SARS-CoV-2 infection: integrative concepts of pathophysiology and case reports. J Neuroinflammation 17, 231 (2020).

20. A. Siderowf et al., Impaired olfaction and other prodromal features in the Parkinson At-Risk Syndrome Study. Mov Disord 27, 406–412 (2012).

21. M. E. Cohen et al., A case of probable Parkinson’s disease after SARS-CoV-2 infection. Lancet Neurol 19, 804–805 (2020).

22. A. Mendez-Guerrero et al., Acute hypokinetic-rigid syndrome following SARS-CoV-2 infection. Neurology 95, e2109–e2118 (2020).

23. I. Faber et al., Coronavirus Disease 2019 and Parkinsonism: A Non-post-encephalitic Case. Mov Disord 35, 1721–1722 (2020).

24. A. Pavel, D. K. Murray, A. J. Stoessl, COVID-19 and selective vulnerability to Parkinson’s disease. Lancet Neurol 19, 719 (2020).

25. P. Brundin, A. Nath, J. D. Beckham, Is COVID-19 a Perfect Storm for Parkinson’s Disease? Trends Neurosci 43, 931–933 (2020).

26. K. J. Ryan et al., A human microglia-like cellular model for assessing the effects of neurodegenerative disease gene variants. Sci Transl Med 9 (2017).

27. B. Korber et al., Tracking Changes in SARS-CoV-2 Spike: Evidence that D614G Increases Infectivity of the COVID-19 Virus. Cell 182, 812–827 e819 (2020).

28. J. Yang et al., Molecular interaction and inhibition of SARS-CoV-2 binding to the ACE2 receptor. Nat Commun 11, 4541 (2020).

29. R. Chen et al., The Spatial and Cell-Type Distribution of SARS-CoV-2 Receptor ACE2 in the Human and Mouse Brains. Front Neurol 11, 573095 (2020).

30. L. Schimmel et al., Endothelial cells are not productively infected by SARS-CoV-2. Clin Transl Immunology 10, e1350 (2021).

31. N. D’Onofrio et al., Glycated ACE2 receptor in diabetes: open door for SARS-COV-2 entry in cardiomyocyte. Cardiovasc Diabetol 20, 99 (2021).

32. C. Conceicao et al., The SARS-CoV-2 Spike protein has a broad tropism for mammalian ACE2 proteins. PLoS Biol 18, e3001016 (2020).

33. L. Yang et al., A Human Pluripotent Stem Cell-based Platform to Study SARS-CoV-2 Tropism and Model Virus Infection in Human Cells and Organoids. Cell Stem Cell 27, 125–136 e127 (2020).

34. A. A. Amarilla et al., A versatile reverse genetics platform for SARS-CoV-2 and other positive-strand RNA viruses. Nat Commun 12, 3431 (2021).

35. S. S. Ousman, P. Kubes, Immune surveillance in the central nervous system. Nat Neurosci 15, 1096–1101 (2012).

36. D. Watterson et al., Preclinical development of a molecular clamp-stabilised subunit vaccine for severe acute respiratory syndrome coronavirus 2. Clin Transl Immunology 10, e1269 (2021).

37. L. Wang, Y. Xiang, Spike Glycoprotein-Mediated Entry of SARS Coronaviruses. Viruses 12 (2020).

38. Y. Chen et al., ACE2-targeting monoclonal antibody as potent and broad-spectrum coronavirus blocker. Signal Transduct Target Ther 6, 315 (2021).

39. A. C. Ferreira et al., SARS-CoV-2 engages inflammasome and pyroptosis in human primary monocytes. Cell Death Discov 7, 43 (2021).

40. S. J. Theobald et al., Long-lived macrophage reprogramming drives spike protein-mediated inflammasome activation in COVID-19. EMBO Mol Med 13, e14150 (2021).

41. J. C. de Rivero Vaccari, W. D. Dietrich, R. W. Keane, J. P. de Rivero Vaccari, The Inflammasome in Times of COVID-19. Front Immunol 11, 583373 (2020).

42. A. Kroemer et al., Inflammasome activation and pyroptosis in lymphopenic liver patients with COVID-19. J Hepatol 73, 1258–1262 (2020).

43. T. S. Rodrigues et al., Inflammasomes are activated in response to SARS-CoV-2 infection and are associated with COVID-19 severity in patients. J Exp Med 218 (2021).

44. C. Junqueira et al., SARS-CoV-2 infects blood monocytes to activate NLRP3 and AIM2 inflammasomes, pyroptosis and cytokine release. medRxiv 10.1101/2021.03.06.21252796 (2021).

45. V. F. Cama et al., The microglial NLRP3 inflammasome is involved in human SARS-CoV-2 cerebral pathogenicity: A report of three post-mortem cases. J Neuroimmunol 361, 577728 (2021).

46. C. B. Jackson, M. Farzan, B. Chen, H. Choe, Mechanisms of SARS-CoV-2 entry into cells. Nat Rev Mol Cell Biol 10.1038/s41580-021-00418-x (2021).

47. J. Shang et al., Cell entry mechanisms of SARS-CoV-2. Proc Natl Acad Sci U S A 117, 11727–11734 (2020).

48. M. A. Niles et al., Macrophages and Dendritic Cells Are Not the Major Source of Pro-Inflammatory Cytokines Upon SARS-CoV-2 Infection. Front Immunol 12, 647824 (2021).

49. M. Z. Ratajczak et al., SARS-CoV-2 Entry Receptor ACE2 Is Expressed on Very Small CD45(-) Precursors of Hematopoietic and Endothelial Cells and in Response to Virus Spike Protein Activates the Nlrp3 Inflammasome. Stem Cell Rev Rep 17, 266–277 (2021).

50. I. Y. Chen, M. Moriyama, M. F. Chang, T. Ichinohe, Severe Acute Respiratory Syndrome Coronavirus Viroporin 3a Activates the NLRP3 Inflammasome. Front Microbiol 10, 50 (2019).

51. J. L. Nieto-Torres et al., Severe acute respiratory syndrome coronavirus E protein transports calcium ions and activates the NLRP3 inflammasome. Virology 485, 330–339 (2015).

52. K. L. Siu et al., Severe acute respiratory syndrome coronavirus ORF3a protein activates the NLRP3 inflammasome by promoting TRAF3-dependent ubiquitination of ASC. FASEB J 33, 8865–8877 (2019).

53. P. Pan et al., SARS-CoV-2 N protein promotes NLRP3 inflammasome activation to induce hyperinflammation. Nat Commun 12, 4664 (2021).

54. F. G. Bauernfeind et al., Cutting edge: NF-kappaB activating pattern recognition and cytokine receptors license NLRP3 inflammasome activation by regulating NLRP3 expression. J Immunol 183, 787–791 (2009).

55. S. F. Dosch, S. D. Mahajan, A. R. Collins, SARS coronavirus spike protein-induced innate immune response occurs via activation of the NF-kappaB pathway in human monocyte macrophages in vitro. Virus Res 142, 19–27 (2009).

56. S. Khan et al., SARS-CoV-2 spike protein induces inflammation via TLR2-dependent activation of the NF-kappaB pathway. bioRxiv 10.1101/2021.03.16.435700 (2021).

57. M. B. Fares, S. Jagannath, H. A. Lashuel, Reverse engineering Lewy bodies: how far have we come and how far can we go? Nat Rev Neurosci 22, 111–131 (2021).

58. A. Filatov, P. Sharma, F. Hindi, P. S. Espinosa, Neurological Complications of Coronavirus Disease (COVID-19): Encephalopathy. Cureus 12, e7352 (2020).

59. P. S. Espinosa, Z. Rizvi, P. Sharma, F. Hindi, A. Filatov, Neurological Complications of Coronavirus Disease (COVID-19): Encephalopathy, MRI Brain and Cerebrospinal Fluid Findings: Case 2. Cureus 12, e7930 (2020).

60. N. Limphaibool, P. Iwanowski, M. J. V. Holstad, D. Kobylarek, W. Kozubski, Infectious Etiologies of Parkinsonism: Pathomechanisms and Clinical Implications. Front Neurol 10, 652 (2019).

61. E. L. Beatman et al., Alpha-Synuclein Expression Restricts RNA Viral Infections in the Brain. J Virol 90, 2767–2782 (2015).

62. C. T. Tulisiak, G. Mercado, W. Peelaerts, L. Brundin, P. Brundin, Can infections trigger alpha-synucleinopathies? Prog Mol Biol Transl Sci 168, 299–322 (2019).

63. V. A. Blanco-Palmero et al., Serum and CSF alpha-synuclein levels do not change in COVID-19 patients with neurological symptoms. J Neurol 10.1007/s00415-021-10444-6 (2021).

64. M. A. Erickson, E. M. Rhea, R. C. Knopp, W. A. Banks, Interactions of SARS-CoV-2 with the Blood-Brain Barrier. Int J Mol Sci 22 (2021).

65. Y. Wu et al., Nervous system involvement after infection with COVID-19 and other coronaviruses. Brain Behav Immun 87, 18–22 (2020).

66. T. E. Poloni et al., COVID-19-related neuropathology and microglial activation in elderly with and without dementia. Brain Pathol 10.1111/bpa.12997, e12997 (2021).

67. E. Song et al., Neuroinvasion of SARS-CoV-2 in human and mouse brain. J Exp Med 218 (2021).

68. P. Kumari et al., Neuroinvasion and Encephalitis Following Intranasal Inoculation of SARS-CoV-2 in K18-hACE2 Mice. Viruses 13 (2021).

69. K. H. Dinnon, 3rd et al., A mouse-adapted model of SARS-CoV-2 to test COVID-19 countermeasures. Nature 586, 560–566 (2020).

70. Ingrid H.C.H.M. Philippens et al., SARS-CoV-2 causes brain inflammation and induces Lewy body formation in macaques. bioRxiv. doi.org/10.1101/2021.02.23.432474 (Preprint posted February 23, 2021) (2021).

71. M. Schwabenland et al., Deep spatial profiling of human COVID-19 brains reveals neuroinflammation with distinct microanatomical microglia-T-cell interactions. Immunity 54, 1594–1610 e1511 (2021).

72. E. G. Brown et al., The Effect of the COVID-19 Pandemic on People with Parkinson’s Disease. J Parkinsons Dis 10, 1365–1377 (2020).

73. R. Cilia et al., Effects of COVID-19 on Parkinson’s Disease Clinical Features: A Community-Based Case-Control Study. Mov Disord 35, 1287–1292 (2020).

74. A. Mullard, NLRP3 inhibitors stoke anti-inflammatory ambitions. Nat Rev Drug Discov 18, 405–407 (2019).

75. X. X. Li, R. J. Clark, T. M. Woodruff, C5aR2 Activation Broadly Modulates the Signaling and Function of Primary Human Macrophages. J Immunol 205, 1102–1112 (2020).

76. V. Deora, E. A. Albornoz, K. Zhu, T. M. Woodruff, R. Gordon, The Ketone Body beta-Hydroxybutyrate Does Not Inhibit Synuclein Mediated Inflammasome Activation in Microglia. J Neuroimmune Pharmacol 12, 568–574 (2017).

77. N. D. Grubaugh et al., An amplicon-based sequencing framework for accurately measuring intrahost virus diversity using PrimalSeq and iVar. Genome Biol 20, 8 (2019).

78. A. A. Amarilla et al., An Optimized High-Throughput Immuno-Plaque Assay for SARS-CoV-2. Front Microbiol 12, 625136 (2021).

79. D. o. V. D. CDC - National Center for Immunization and Respiratory Diseases (NCIRD) (2021) CDC 2019-nCoV Real-Time RT-PCR Diagnostic Panel. p Acceptable Alternative Primer and Probe Sets

80. A. Isaacs et al., Combinatorial F-G Immunogens as Nipah and Respiratory Syncytial Virus Vaccine Candidates. Viruses 13 (2021).

81. J. ter Meulen et al., Human monoclonal antibody combination against SARS coronavirus: synergy and coverage of escape mutants. PLoS Med 3, e237 (2006).

