## Supplementary material for "SARS-CoV-2 drives NLRP3 inflammasome activation in human microglia through spike-ACE2 receptor interaction": Suplemmentary figures

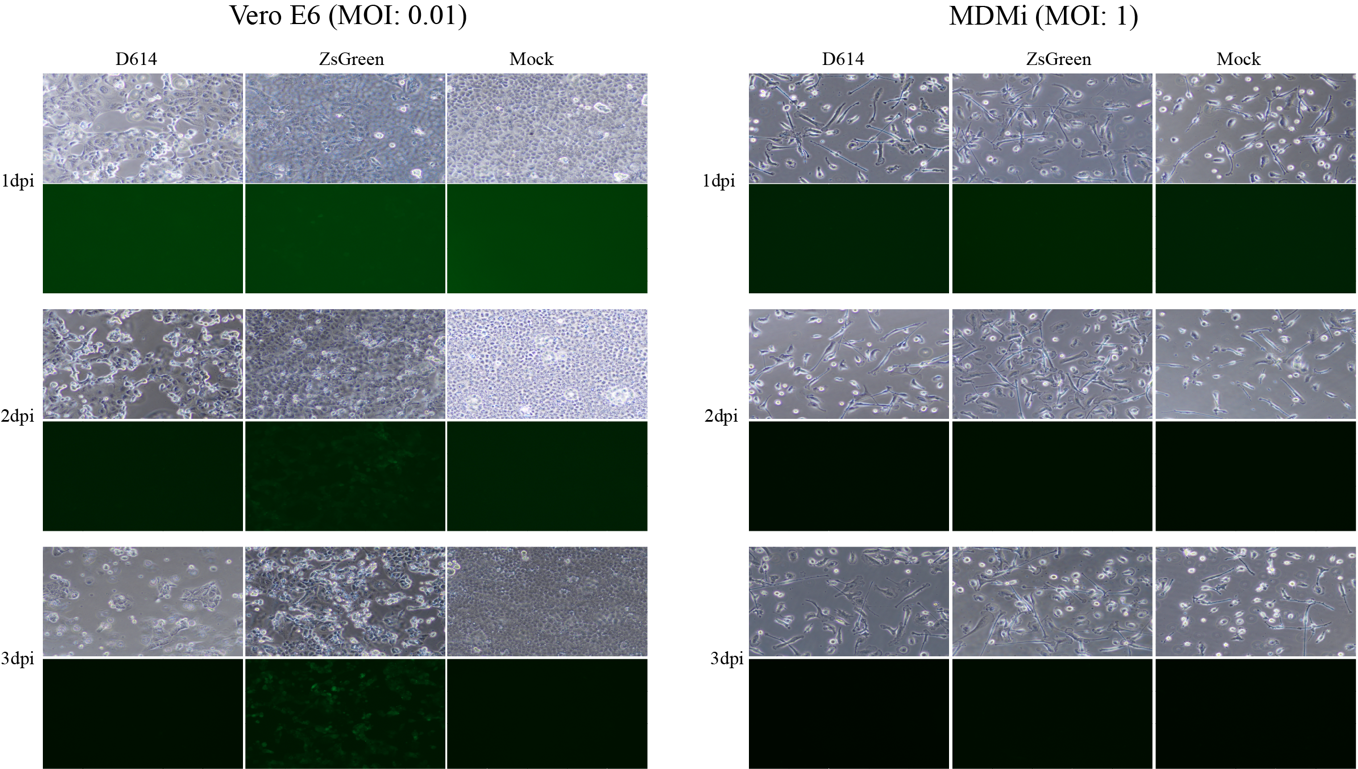

**Supplementary Figure 1.** SARS-CoV-2 replication on MDMi (at MOI of 1) and Vero E6 (at MOI of 0.01) using SARS-CoV-2 reporter virus expressing ZsGreen fluorescent protein compared to WT virus (D614) assessed directly under microscopy. Images on the left and right panel are from infected Vero E6 and MDMi cell, respectively. Images for each time point were taken under bright-field (on top) and with the fluorescence filter (on bottom) using 40x magnification.

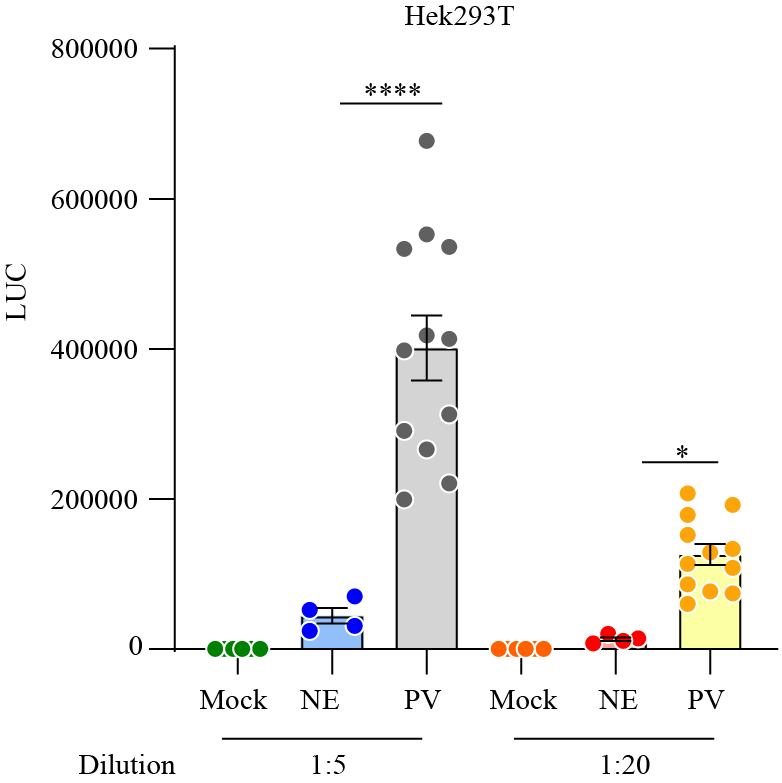

**Supplementary Figure 2.** Validation of Intracellular luciferase level (LUC) delivered by pseudo-virus (PV) particle for SARS-CoV-2 in HEK293T compared to the non-glycoprotein control (NE). Data are means ± SD from one experiment. *P < 0.05, **P < 0.01, and ***P < 0.001 and **** P < 0.0001 by one-way analysis of variance (ANOVA) with Tukey’s post hoc test.

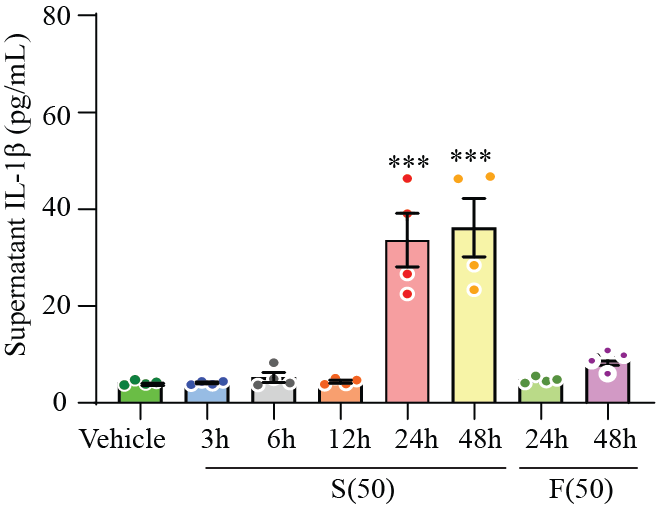

**Supplementary Figure 3.** Time course of IL-1β secretion in the supernatant of MDMi exposed to S-Clamp (S;50 μg) of F-clamp (F;50 μg). Data are means ± SEM from at least three different donors. ***P < 0.001 by one-way analysis of variance (ANOVA) with Tukey’s post hoc test.
